# MSLipidMapper: a pathway-centered lipidome analysis environment linking lipid class, acyl-chain subsets, and multi-omics data

**DOI:** 10.64898/2026.05.21.726751

**Authors:** Takaki Oka, Kozo Nishida, Takeshi Harayama, Hiroshi Tsugawa

## Abstract

Lipids exhibit extensive structural diversity arising from variation in lipid classes, subclasses, and acyl-chain compositions, making systematic interpretation of lipidomics data challenging. Although untargeted lipidomics enables the quantification of hundreds to thousands of lipid molecular species, downstream analyses often treat pathway-level summaries, molecular-species visualization, structural subsetting, and multi-omics interpretation as separate steps. Here, we present MSLipidMapper, an R/Shiny-based lipidomics data exploration environment for pathway-centered and structure-aware analysis of annotated lipidomics datasets. MSLipidMapper reconstructs annotated lipid peak tables as Bioconductor SummarizedExperiment objects, thereby organizing quantitative lipid abundance values, sample metadata, lipid subclass annotations, and parsed acyl-chain features within a unified data structure. Lipid molecular species are summarized on static, curated lipid metabolic pathway maps at the subclass level while retaining direct links to the underlying molecular species and acyl-chain annotations. This design enables users to inspect molecular-species patterns underlying each pathway node, define lipid subsets based on structural features such as specific acyl chains, and re-project these subsets onto the same pathway context. Gene or protein expression data can also be overlaid on pathway-associated reactions to support multi-layer interpretation of lipid metabolism. The program is showcased using publicly available aging lipidome datasets of mice, illustrating how subclass-level pathway summaries can be connected to molecular-species heatmaps, acyl-chain-defined subsets, and transcriptome or proteome information.

## INTRODUCTION

Untargeted lipidomics enables the quantitative profiling of hundreds to thousands of structurally diverse lipid molecules within a single experiment^1^. This approach has provided important opportunities to investigate lipid remodeling across biological processes and disease contexts, including aging, metabolic disorders, cardiovascular disease, cancer, and inflammation^2–4^. Unlike many small-molecule metabolites, lipids are organized into hierarchical molecular families defined by lipid classes, subclasses, acyl-chain compositions, degrees of unsaturation, and other structural features. This organization is reflected in classification systems such as LIPID MAPS and is biologically meaningful because lipid metabolism is mediated by enzymatic pathways that synthesize, remodel, and degrade lipid molecules in a class- and acyl-chain-dependent manner^6^. Therefore, biological interpretation of lipidomics data requires analytical frameworks that preserve the relationships among lipid subclasses, molecular species, and structural features while enabling systematic exploration of lipid abundance patterns.

A typical lipidomics workflow consists of spectral processing, lipid annotation, and downstream biological interpretation^7^. The development of open-source software such as MS-DIAL has facilitated vendor-independent peak detection, quantification, and lipid annotation, enabling researchers to obtain annotated lipid peak tables from liquid chromatography–mass spectrometry (LC–MS) experiments in a standardized manner^8^. In contrast, downstream lipidomics analysis remains less standardized and is often performed using custom R or Python scripts together with multiple visualization and web-based tools^9^. As a result, statistical analysis, lipid-class summarization, molecular-species visualization, pathway-level interpretation, and integration with transcriptomic or proteomic data are frequently conducted as separate steps. This fragmentation makes it difficult to reproducibly connect lipid subclass-level abundance changes with the molecular species and structural features that underlie them.

Several computational platforms have been developed to support downstream analysis of metabolomics and lipidomics data. General metabolomics platforms, such as MetaboAnalyst and MetaboAnalystR, provide comprehensive workflows for statistical analysis, visualization, enrichment analysis, pathway analysis, and multi-omics integration^10,11^. These tools are widely used for interpreting metabolite-level changes in biological pathways. However, because they are designed for broad metabolomics applications, lipid molecules are generally handled as individual metabolite features rather than as hierarchical molecular families defined by lipid subclasses and acyl-chain compositions.

Lipidomics-oriented tools address some of these lipid-specific requirements. The R/Bioconductor package “lipidr” supports data inspection, normalization, univariate and multivariate analysis, and visualization of lipidomics datasets^12^. LipidSig extends lipidomics data mining to enrichment, correlation, machine learning, and network analysis^13^. Network-based tools such as LINEX2 further represent biochemical relationships among lipid species, subclasses, and acyl-chain transformations, enabling detailed exploration of lipid remodeling networks^14^. These tools provide valuable functionality for lipidomics data analysis, but they are not primarily designed to summarize lipid abundance changes on fixed, curated lipid metabolic pathway maps while preserving direct links to the molecular species and structural subsets underlying each pathway-level summary. Because untargeted lipidomics datasets often contain hundreds to thousands of annotated lipid species, molecular-species-level networks can become large and visually complex. This complexity can make it difficult to interpret lipid remodeling within a stable pathway context and to trace structurally defined lipid groups, such as lipids containing a specific acyl chain, across related lipid subclasses. There is therefore a need for a pathway-centered framework that summarizes lipidomics data on static, curated lipid metabolic maps while retaining access to molecular-species-level and acyl-chain-resolved information.

To address this need, we developed MSLipidMapper, an R/Shiny-based lipidomics data exploration environment for pathway-centered and structure-aware analysis of LC–MS lipidomics datasets. MSLipidMapper reconstructs annotated lipid peak tables as Bioconductor SummarizedExperiment objects, in which quantitative abundance data, sample metadata, lipid subclass annotations, and parsed structural features are maintained within a unified data structure. Using this representation, lipid molecular species can be summarized at the subclass level and projected onto static, curated lipid metabolic pathway maps. At the same time, the links between subclass-level summaries and their underlying molecular species are preserved, allowing users to inspect which lipid species and acyl-chain compositions contribute to each pathway-level abundance pattern. MSLipidMapper provides an integrated workflow for statistical analysis, molecular-species visualization, pathway visualization, structure-defined subsetting, and multi-omics integration. In the pathway view, lipid abundance values are summarized on lipid subclass nodes, enabling users to interpret lipidomics data within a consistent pathway context. Because each subclass-level summary retains links to the corresponding molecular species, users can examine whether the observed abundance pattern is broadly shared across the subclass or driven by specific molecular species. Lipid groups can also be redefined using parsed structural features, such as the presence of specific acyl chains, enabling acyl-chain-defined subsets to be re-projected onto the same curated pathway map. In addition, gene or protein expression data can be overlaid onto pathway-associated reactions, supporting multi-layer interpretation of lipid metabolism. In this study, we describe the design and implementation of MSLipidMapper and demonstrate its utility using publicly available aging lipidome datasets of mice. These examples illustrate how subclass-level pathway summaries can be connected to molecular-species heatmaps, acyl-chain-defined subsets, and transcriptomic information within a single analytical environment. MSLipidMapper provides an interpretable and extensible framework for reproducible downstream lipidomics analysis and supports pathway-centered exploration of structure-dependent lipid abundance patterns in complex lipidomics datasets.

## EXPERIMENTAL SECTION

### Data model and representation

MSLipidMapper represents lipidomics datasets using the Bioconductor “SummarizedExperiment” framework^15^, in which quantitative lipid abundance values, feature-level annotations, and sample metadata are stored in a unified container. Quantitative abundance matrices are stored as assays, sample-level metadata are stored as “colData”, and lipid annotations are stored as “rowData”. Each lipid molecular species is treated as a feature and is associated with structured annotations, including lipid subclass, parsed acyl/alkyl-chain composition, and derived structural attributes such as total chain length and degree of unsaturation. This data structure enables downstream analyses to be performed consistently across sample-level, subclass-level, molecular-species-level, and structure-defined subset representations.

### Parsing of lipid names and extraction of structural features

MSLipidMapper uses lipid names assigned by upstream annotation tools as input and extracts the structural features required for downstream analysis. For lipid names generated by MS-DIAL^8^, acyl/alkyl-chain information is extracted using predefined rule-based parsing procedures (Supplementary Table 1). For lipid tables generated by other annotation tools, lipid names can be parsed using Goslin^16^. Parsed structural features, including lipid subclass, acyl/alkyl-chain composition, chain length, and degree of unsaturation, are stored in the feature metadata of the SummarizedExperiment object. Chain descriptors are defined by carbon number and degree of unsaturation, such as 16:0 and 20:4. These annotations are then used for filtering, grouping, aggregation, and pathway projection of lipid species based on user-defined structural criteria.

### Mapping of lipid species to pathway nodes

Lipid molecular species are mapped to pathway nodes using predefined mapping keys in curated lipid metabolic pathway templates. These keys are based on MS-DIAL lipid class/subclass names or LIPID MAPS^17^ identifiers, depending on the pathway template. Input lipid names and associated identifiers are matched to these keys to assign each molecular species to the corresponding pathway node. Molecular species assigned to the same node are grouped, and their quantitative values are aggregated to generate node-level summaries. The mapping table is retained to preserve the correspondence between pathway nodes and the underlying lipid molecular species.

### Pathway projection and structure-defined subset analysis

Quantitative lipid abundance values are aggregated for each pathway node according to the mapping between lipid molecular species and pathway nodes. For global subclass-level visualization, all molecular species assigned to a given node are used to calculate node-level summaries. For structure-defined subset visualization, lipid molecular species are first filtered using parsed structural features, such as the presence of a specific acyl or alkyl chain, and the filtered species are then aggregated using the same node mapping. The resulting subset-level values are displayed on the same curated pathway template as the global subclass-level summaries. This design enables direct comparison between overall subclass-level abundance patterns and structure-defined subset patterns within a consistent pathway context.

### Multi-omics integration

Gene and protein expression data are integrated by mapping genes or proteins to enzyme-associated reaction edges in the curated lipid metabolic pathway templates. Input expression tables are required to contain Ensembl identifiers, which are used as mapping keys between user-provided omics data and pathway-associated genes. Quantitative expression values are imported as gene- or protein-level abundance tables and visualized alongside lipid abundance summaries within the pathway view.

### Construction of lipid metabolic pathway templates

The lipid metabolic pathway templates used in MSLipidMapper were constructed as mammalian lipid metabolic pathway maps by integrating information from publicly available lipid resources, including LIPID MAPS, together with representative literature describing lipid metabolic pathways^17,18^. The templates use an abstracted pathway representation in which lipid subclasses are represented as nodes and biochemical relationships between subclasses are represented as edges. This representation enables lipid molecular species to be summarized at the subclass level on pathway maps while retaining links to the underlying molecular species. Enzyme– gene relationships were manually curated from the literature and incorporated into the pathway templates to support multi-omics visualization. The resulting pathway templates are implemented as network objects that can be visualized and edited within the application. Users can also import custom pathway structures in standard network formats, including .cyjs and .gml.

### Implementation and visualization architecture

MSLipidMapper is implemented in R as a modular Shiny application, in which individual analytical components operate on a shared SummarizedExperiment object. MSLipidMapper is implemented in R as a modular Shiny application. Individual analytical components operate on a shared SummarizedExperiment object, allowing lipid abundance values, feature annotations, sample metadata, and optional multi-omics data to be accessed consistently across the workflow. Pathway visualization is performed using Cytoscape.js^19^ embedded within the Shiny interface. Communication between the R backend and the browser-based frontend is mediated by a Plumber-based HTTP API^20^. In this architecture, quantitative results and node-associated visual assets, including heatmaps and summary plots, are generated in R and exposed as HTTP-accessible resources through the Plumber endpoint. Cytoscape.js retrieves these resources through HTTP requests and renders them within the pathway view. Upon user interaction, such as pathway-node selection or application of structural filters, the R backend updates the corresponding outputs and exposes the updated resources through the same API.

## RESULTS

### Overview of a static pathway-centered framework for hierarchical lipidomics analysis

To support systematic interpretation of complex lipidomics datasets, we developed MSLipidMapper, a pathway-centered analysis framework that summarizes lipidomics data on curated lipid metabolic maps while preserving molecular-species-level and acyl-chain-resolved information. MSLipidMapper accepts annotated lipid peak tables generated by tools such as MS-DIAL as input and reconstructs them as Bioconductor SummarizedExperiment objects, in which quantitative measurements, sample metadata, lipid subclass annotations, and parsed structural features are maintained within a unified data structure.

MSLipidMapper organizes downstream lipidomics analysis using a hierarchical representation of lipid molecules (**Figure 1**). Individual lipid molecular species are first assigned to lipid subclasses, which serve as the primary units for mapping onto static, curated metabolic pathway templates. This design allows abundance values from many lipid molecular species to be summarized on pathway nodes without expanding the visualization into a large molecular-species-level network. At the same time, the correspondence between each pathway-node summary and its underlying molecular species is retained, enabling users to inspect which lipid species and structural features contribute to the summarized abundance pattern. Because lipid subclass assignments and acyl-chain annotations are stored as structured feature metadata, molecular species can be aggregated not only by subclass but also according to user-defined structural criteria. For example, lipid species containing a specific acyl chain can be selected from the same underlying dataset and summarized on the same pathway template. This enables global subclass-level summaries and structure-defined subset summaries to be compared within a fixed pathway context without manual restructuring of the input data. By maintaining these relationships within a shared data structure, MSLipidMapper establishes a reproducible link between subclass-level pathway summaries, molecular-species-level measurements, and acyl-chain annotations. This framework allows users to examine whether a pathway-node abundance pattern reflects broad changes across a lipid subclass or is driven by specific molecular species or acyl-chain-defined subsets.

**Figure 1.**
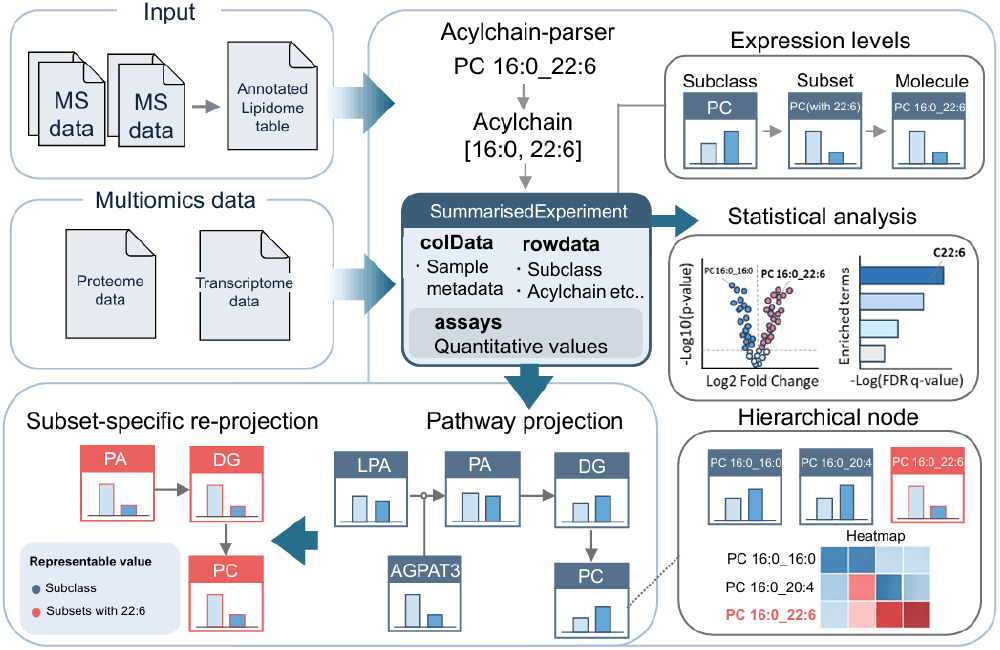
Overview of the MSLipidMapper framework for static pathway-centered lipidomics analysis. MSLipidMapper integrates lipidomics data, pathway visualization, structure-aware exploration, and multi-omics information within a unified analytical framework. Annotated lipid tables derived from mass spectrometry data are processed to extract lipid subclass and acyl-chain information and are reconstructed as Bioconductor SummarizedExperiment objects containing quantitative abundance values (assays), sample metadata (colData), and lipid structural annotations (rowData). Lipid molecular species are organized hierarchically into subclasses, individual molecular species, and structure-defined subsets, allowing abundance values to be summarized and visualized on static, curated lipid metabolic pathway maps. Links between pathway nodes and the underlying molecular species are retained, enabling molecular-species-level inspection, acyl-chain-defined subset analysis, and subset-specific pathway re-projection. Transcriptomic and proteomic data can also be incorporated to support multi-layer interpretation of lipid metabolism.

### Pathway-centered visualization links lipid-subclass summaries to molecular-species patterns

A central feature of MSLipidMapper is its ability to connect pathway-level lipid subclass summaries with molecular-species-level abundance patterns within a static, curated pathway context. In the pathway view, each node represents a lipid subclass and displays its abundance distribution across samples, with individual points representing samples and horizontal lines indicating group medians. This representation allows lipidomics datasets containing many molecular species to be summarized on interpretable pathway maps without expanding the visualization into a large molecular-species-level network. Each pathway node is linked to the molecular species assigned to the corresponding lipid subclass through the shared SummarizedExperiment object. When a user selects a pathway node, the associated molecular species are retrieved and visualized as a heatmap, enabling inspection of the abundance patterns underlying the subclass-level summary. This node-linked visualization allows users to assess whether a subclass-level abundance pattern reflects broad changes across many lipid species or is driven by a restricted subset of molecules.

The same data representation also enables molecular species to be selected according to parsed structural attributes, such as the presence of specific acyl or alkyl chains. Selected lipid species can then be regrouped by subclass and projected onto the same curated pathway template used for global subclass-level visualization. This functionality allows users to compare overall subclass-level abundance patterns with structure-defined subset patterns, such as lipids containing a specific acyl chain, within a consistent pathway context.

Compared with existing metabolomics and lipidomics tools, MSLipidMapper is characterized by the combination of static curated pathway visualization, subclass-level node summarization, molecular-species-level inspection, acyl-chain-defined subset tracking, and multi-omics overlay within a single workflow (**Table 1**). This hierarchical exploration is examplified by using phosphatidylinositol, in which a subclass-level pathway node is linked to molecular-species heatmaps and individual lipid abundance plots (**Figure 2**). By linking subclass-level pathway summaries, molecular-species heatmaps, individual lipid abundance plots, and acyl-chain-defined subset projections, MSLipidMapper provides an integrated framework for tracing lipid abundance patterns across multiple levels of lipid organization.

**Table 1.**
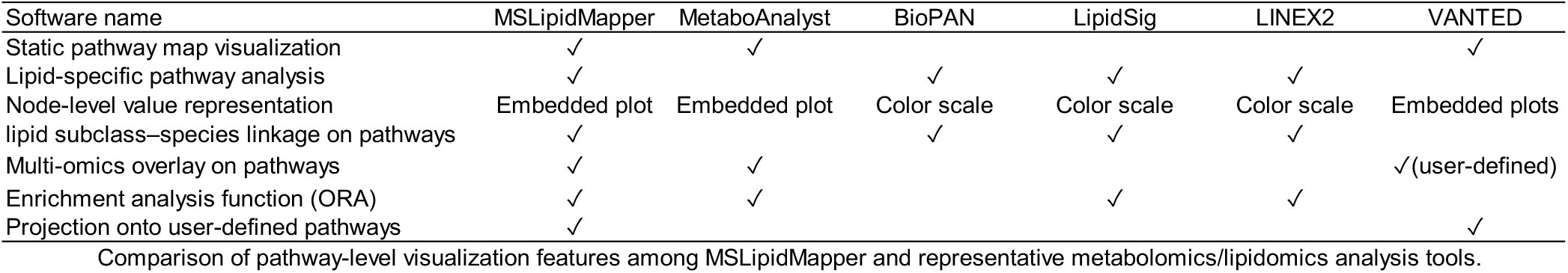
Pathway visualization feature in metabolomics tools.

Comparison of pathway-level visualization features among MSLipidMapper and representative metabolomics/lipidomics analysis tools.
| Software name | MSLipidMapper | MetaboAnalyst | BioPAN | LipidSig | LINEX2 | VANTED |
| --- | --- | --- | --- | --- | --- | --- |
| Static pathway map visualization | ✓ | ✓ |  |  |  | ✓ |
| Lipid-specific pathway analysis | ✓ |  | ✓ | ✓ | ✓ |  |
| Node-level value representation | Embedded plot | Embedded plot | Color scale | Color scale | Color scale | Embedded plots |
| lipid subclass-species linkage on pathways | ✓ |  | ✓ | ✓ | ✓ |  |
| Multi-omics overlay on pathways | ✓ | ✓ |  |  |  | ✓(user-defined) |
| Enrichment analysis function (ORA) | ✓ | ✓ |  | ✓ | ✓ |  |
| Projection onto user-defined pathways | ✓ |  |  |  |  | ✓ |

**Figure 2.**
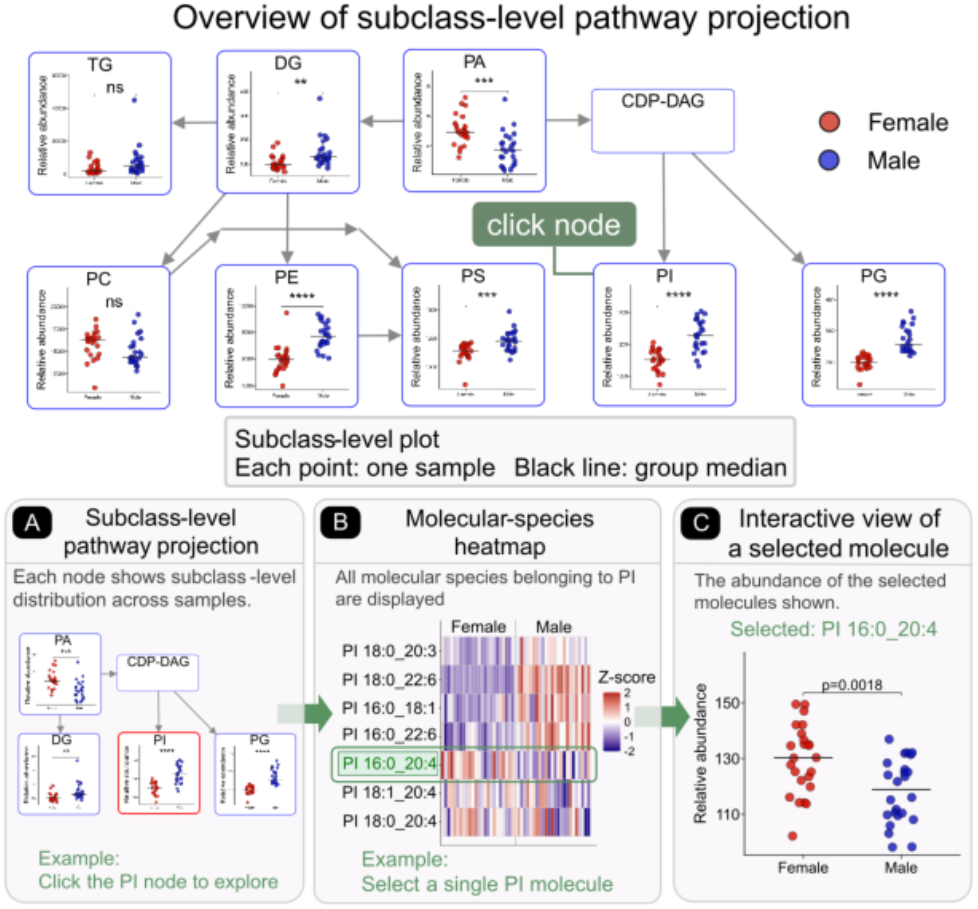
Hierarchical exploration of lipidomics data in MSLipidMapper. Sex-dependent differences in kidney lipid profiles are shown as an example. Pathway projection at subclass and molecular species levels shows that some molecular species of phosphatidylinositol (PI) display distinct trends as compared to global PI levels. (a) Subclass-level pathway visualization, in which each pathway node represents a lipid subclass and displays its abundance distribution across samples. Individual points indicate samples, and horizontal lines indicate group medians. (b) Molecular-species heatmap linked to a selected subclass node. Selecting a node, such as phosphatidylinositol (PI), retrieves the corresponding molecular species from the shared data object and displays their abundance patterns, with rows representing molecular species and columns representing samples. (c) Molecular-species-level inspection of a selected lipid species. Selecting a lipid species from the heatmap, such as PI 16:0_20:4, displays its abundance distribution across samples.

### Reanalysis of kidney lipidomics illustrates acyl-chain-dependent sphingolipid patterns masked by subclass-level summaries

To demonstrate the utility of MSLipidMapper for pathway-centered interpretation of lipidomics data, we reanalyzed kidney lipidomics and transcriptomics data from a public aging mouse study^21^. Projection of lipid subclass-level summaries onto curated sphingolipid metabolic pathways recapitulated the male-biased accumulation of hexosylceramide (HexCer) (P=1.1 × 10^−17^) (**Figure 3a**). This subclass-level increase was accompanied by higher expression of Ugt8a in males than in females (P=4.3 × 10^−16^), consistent with the role of UGT8 in glycosphingolipid biosynthesis. Although this subclass-level pathway summary provided an overview of sex-dependent sphingolipid abundance patterns in kidney, it did not indicate whether all HexCer molecular species contributed uniformly to the male-specific HexCer profile. We therefore inspected the molecular-species-level abundance patterns underlying the HexCer node using the node-linked heatmap. Most HexCer species showed higher abundance in males; however, a subset of species exhibited distinct patterns. In particular, HexCer 18:1;2O/20:0 showed higher abundance in females than in males (P = 0.00316) (**Figure 3b and 3c**), indicating that the global HexCer summary masked species-level heterogeneity.

**Figure 3.**
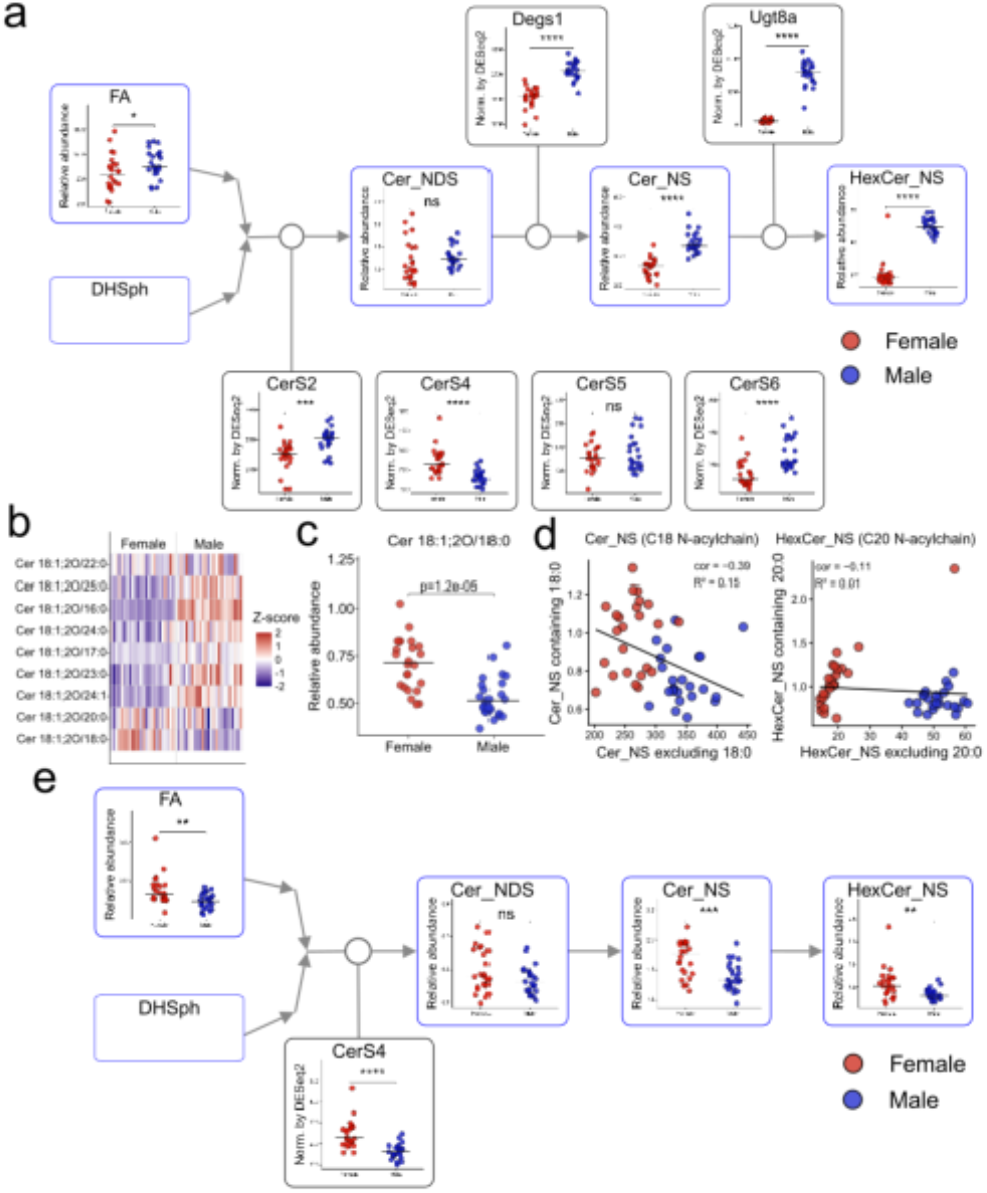
Acyl-chain-resolved analysis of sphingolipid abundance patterns in kidney lipidomics data. (a) Subclass-level pathway projection of sphingolipid metabolism. Each pathway node represents the abundance distribution of a lipid subclass; individual points indicate samples, and black horizontal lines indicate group medians. Gene expression levels are mapped onto circular nodes positioned on reaction edges. (b) Molecular-species heatmap of HexCer_NS. Rows represent lipid molecular species, columns represent samples, and values are shown as Z-scores. (c) Abundance distribution of HexCer 18:1;2O/20:0 selected from the heatmap. Each point represents a sample, and black horizontal lines indicate group medians. (d) Correlation between acyl-chain-defined subsets and the remaining species within the same subclass. The left panel shows HexCer_NS species containing 20:0 acyl chains, and the right panel shows Cer_NS species containing 18:0 acyl chains. The x-axis represents species excluding the specified acyl chain, and the y-axis represents the corresponding acyl-chain-defined subset. Pearson’s correlation coefficient (r) and coefficient of determination (R^2^) are shown. (e) Subset-specific pathway projection of sphingolipid species containing 18:0 or 20:0 acyl chains. Each node represents the abundance distribution calculated from the selected subset, and gene expression levels are mapped onto circular nodes on reaction edges. P values in panel (c) were calculated using two-sided Wilcoxon rank-sum tests.

Correlation analysis between acyl-chain-defined subsets and the remaining species within the same subclass showed that the 20:0-containing subset of HexCer non-hydroxyfatty acid-sphingosine (NS) was only weakly correlated with the rest of the subclass (r = −0.11, R^2^ = 0.01). Similarly, the 18:0-containing subset of Cer_NS showed reduced correlation with the remaining Cer_NS species (r = −0.39, R^2^ = 0.15) (**Figure 3d**). These results suggest that C18–C20-containing sphingolipid subsets exhibit abundance patterns that differ from the corresponding subclass-level summaries. We then summarized sphingolipid species containing 18:0 or 20:0 acyl chains and projected these subset-defined abundance patterns onto the same curated pathway map. In contrast to the male-biased HexCer subclass-level pattern, the subset-based analysis showed higher abundance of 18:0- and 20:0-containing sphingolipid species in females (**Figure 3e**). The expression of ceramide synthase 4 (CerS4), which has been reported to exhibit selectivity toward C18–C20 acyl-CoA substrates^22^, was also higher in females (P = 5.8 × 10^−6^). This expression pattern was consistent with the female-biased abundance of C18–C20-containing sphingolipid subsets.

Together, these analyses demonstrate how MSLipidMapper connects subclass-level pathway summaries with molecular-species-level information, acyl-chain-defined subsets, and transcriptomic data within a single analytical environment. The kidney lipidomics example illustrates that pathway-level summaries can be used as an entry point for exploring molecular-species- and acyl-chain-resolved abundance patterns in public lipidomics datasets.

## DISCUSSION

In this study, we developed MSLipidMapper, a pathway-centered lipidomics data exploration environment designed to summarize LC–MS-based lipidomics data on curated lipid metabolic pathway maps while retaining access to molecular-species- and acyl-chain-resolved information. A central concept of MSLipidMapper is to use static pathway templates as an interpretable biological context, rather than expanding lipidomics datasets into large molecular-species-level networks. In this framework, subclass-level abundance summaries displayed on pathway nodes remain linked to the underlying molecular species and parsed structural features. By reconstructing annotated lipid peak tables as Bioconductor SummarizedExperiment^19^ objects, MSLipidMapper maintains quantitative measurements, sample metadata, lipid subclass annotations, and acyl-chain information within a unified data structure. This design enables users to move between pathway-level subclass summaries, molecular-species-level abundance patterns, acyl-chain-defined subsets, and gene or protein expression data related to lipid-metabolic enzymes without manual restructuring of the dataset.

The application to a publicly available aging mouse lipidome dataset^21^ demonstrates how MSLipidMapper can be used to explore lipidomics data across multiple levels of lipid organization. In the kidney dataset, projection of subclass-level lipid abundance onto a curated sphingolipid pathway provided an overview of sex-associated sphingolipid abundance patterns, including male-specific accumulation of HexCer accompanied by higher Ugt8a expression. However, inspection of the molecular species underlying the HexCer node showed that specific HexCer species exhibited abundance patterns that differed from the overall subclass-level summary. In particular, the C20-containing HexCer species showed a pattern distinct from the global HexCer trend. Subsequent acyl-chain-defined subset analysis further illustrated that C18–C20-containing sphingolipid subsets could be traced on the same pathway context and compared with expression patterns of related lipid-metabolic genes, including CerS4. This example highlights the utility of MSLipidMapper for connecting pathway-level summaries to molecular-species- and acyl-chain-resolved information.

A key advantage of MSLipidMapper is that it preserves the relationship between subclass-level pathway summaries, molecular species, and structural annotations. Conventional visualizations such as volcano plots and heatmaps are useful for identifying individual altered lipids, but they often do not show how these molecules relate to lipid metabolic pathways or whether the observed abundance patterns are shared across structurally related lipid groups. Conversely, subclass-level summaries provide an interpretable overview but may obscure molecular-species- or acyl-chain-specific abundance patterns. MSLipidMapper bridges these levels by allowing lipid species to be summarized on curated pathway maps while retaining direct links to the underlying molecular species and parsed structural features. This approach enables users to examine whether a subclass-level abundance pattern reflects broad changes across many molecular species or is driven by a structurally defined subset of lipids.

MSLipidMapper is complementary to existing lipidomics tools rather than a replacement for them. Network-based tools such as LipidSig^13^ and LINEX2^14^ provide powerful approaches for exploring lipid–lipid relationships, biochemical reaction networks, and structural transformations among lipid species. However, when hundreds to thousands of molecular species are represented as network nodes, the resulting visualization can become large and difficult to interpret within a fixed biological pathway context. MSLipidMapper addresses a different need by summarizing lipidomics data on static, curated lipid metabolic maps while preserving links to molecular species and acyl-chain annotations. This design is particularly useful when the goal is to interpret lipid abundance patterns within established metabolic pathways and to follow structurally defined lipid groups, such as lipids containing a specific acyl chain, across related lipid subclasses.

Several limitations should be considered. The accuracy of pathway-level interpretation depends on the quality of lipid annotation and structural parsing. Misannotation or incomplete structural resolution, such as uncertain sn-position, double-bond position, hydroxylation state, or oxidized modification, may affect the interpretation of acyl-chain-defined subsets. In addition, MSLipidMapper summarizes lipid molecular species at the subclass level on curated pathway templates; therefore, it does not replace molecular-species-level network models when detailed reaction-level relationships among individual lipid species are required. Moreover, the curated pathway templates used in MSLipidMapper represent the current state of lipid metabolic knowledge. Because lipid metabolic pathways and enzyme–lipid relationships are continuously updated as new findings emerge, the pathway maps will require periodic revision and curation to remain up to date. The current implementation is intended to support interpretable exploration and hypothesis generation from annotated lipidomics datasets, rather than to infer causal mechanisms of lipid regulation.

In summary, MSLipidMapper provides a unified framework for pathway-centered interpretation of lipidomics data while preserving molecular-species- and acyl-chain-level information. By combining static curated pathway maps with molecular-species-level inspection, acyl-chain-defined subset analysis, and multi-omics integration, MSLipidMapper enables lipid abundance patterns to be traced across subclass, molecular-species, and structural levels. The kidney lipidomics example demonstrates how the program can be applied to public lipidomics datasets to explore structure-dependent lipid abundance patterns within a curated pathway context. This framework should facilitate reproducible and interpretable exploration of complex lipidomics datasets and support the generation of hypotheses about lipid metabolism.

## CODE AVAILABILITY

The source code for MSLipidMapper is available at GitHub: https://github.com/systemsomicslab/MSLipidMapper.

## AUTHOR INFORMATION

### Author Contributions

T.O. and H.T. designed the study. T.O. performed computational data analysis. K.N. and T.H. provided technical advice. H.T. supervised the project. T.O. and H.T. wrote the manuscript. All authors discussed the results and contributed to the final version of the manuscript.

## ACKNOWLEDGMENT

This study represents a portion of the dissertation submitted by Takaki Oka to the Tokyo University of Agriculture and Technology in partial fulfillment of the requirements for his Ph.D. This research was supported by the Japan Science and Technology Agency (JST) BOOST (JPMJBS2420 to T.O.), JST ERATO (JPMJER2101 to H. T.), JST FOREST (JPMJFR230H to H.T.), JST NBDC (JPMJND2305 to H.T.), JST ASPIRE (JPMJAP2505 to H.T.), the JSPS KAKENHI (24K02011, 24H00043, 24H00392, 24K21269, 25H01425, and 25H01426 to H.T.), the Japan Agency for Medical Research and Development (AMED) under Infectious Diseases Research and Infrastructure (JP25wm0325071, H.T.), the French National Research Agency (ANR-25-CE44-1525), and Inserm (IRP AtypicoLipid to T.H. and H.T.).

## REFERENCES

1. Tsugawa, H. et al. A lipidome atlas in MS-DIAL 4. Nat Biotechnol 38, 1159–1163 (2020).

2. Salihovic, S. et al. Recent Advances towards Mass Spectrometry-Based Clinical Lipidomics. Current Opinion in Chemical Biology 123, 9744–9784 (2023).

3. Hornburg, D. et al. Dynamic Lipidome Alterations Associated with Human Health, Disease and Ageing. Nat. Metab. 5, 1578–1594 (2023).

4. Surendran, A., Zhang, H., Stamenkovic, A., Ravandi, Lipidomics and Cardiovascular Disease. Biochim. Biophys. Acta, Mol. Basis Dis. 1871, 5 (2025).

5. Liebisch, G. et al. Update on LIPID MAP. Classification, Nomenclature, and Shorthand Notation for MS-Derived Lipid Structures. J. Lipid Res. 61, 1539–1555 (2020).

6. Harayama, T. & Riezman, H. Nat. Rev. Mol. Cell Biol. 19, 281–296 (2018).

7. Köfeler, H. C. et al. Recommendations for Good Practice in MS-Based Lipidomics. J. Lipid Res. 62, 100138 (2021).

8. Takeda, H. et al. MS-DIAL 5 multimodal mass spectrometry data mining unveils lipidome complexities. Nat Commun 15, 9903 (2024).

9. Best practices and tools in R and Python for statistical processing and visualization of lipidomics and metabolomics data. Nature Communications 16, 8714 (2025).

10. Pang, Z., Lu, Y., Zhou, G., Hui, F., Xu, L., Viau, C., Spigelman, A. F., MacDonald, P. E., Wishart, D. S., Li, S., Xia, J. MetaboAnalyst 6.0: Towards a Unified Platform for Metabolomics Data Processing, Analysis and Interpretation. Nucleic Acids Res. 52 (W1) (2024). pW398–W406. 10.1093/nar/gkae253.

11. Pang, Z., Zhou, G., Ewald, J., Chang, L., Hacariz, O., Basu, N., Xia, J. MetaboAnalystR 4.0: A Unified LC–MS Workflow for Global Metabolomics. Nat. Commun. 15, 3675 (2024).

12. Mohamed, A., Molendijk, J., Hill, M. M. lipidr: A Software Tool for Data Mining and Analysis of Lipidomics Datasets. J. Proteome Res. 19, 2890–2897 (2020).

13. Liu, C.-H. et al. LipidSig 2.0: Integrating Lipid Characteristic Insights into Advanced Lipidomics Data Analysis. Nucleic Acids Res. 52, W390–W397 (2024).

14. Rose, T. D. et al. Lipid Network and Moiety Analysis for Revealing Enzymatic Dysregulation and Mechanistic Alterations from Lipidomics Data. Brief. Bioinform. 24 (1), bbac572 (2023).

15. Huber, W. et al. Orchestrating High-Throughput Genomic Analysis with Bioconductor. Nat. Methods 12, 115–121 (2015).

16. Kopczynski, D., Hojmann, N., Peng, B., Liebisch, G., Spener, F., Ahrends, R. Goslin 2.0 Implements the Recent Lipid Shorthand Nomenclature for MS-Derived Lipid Structures. Anal. Chem. 94 (16), 6097–6101 (2022).

17. Conroy, M. J. et al. LIPID MAPS: Update to Databases and Tools for the Lipidomics Community. Nucleic Acids Res. 52 (D1), D1677–D1682 (2024).

18. Valentine, W. J., Yanagida, K., Kawana, H., Kono, N., Noda, N. N., Aoki, J. Update and Nomenclature Proposal for Mammalian Lysophospholipid Acyltransferases, Which Create Membrane Phospholipid Diversity. J. Biol. Chem. 298 (1), 101470 (2022).

19. Franz, M., Lopes, C. T., Huck, G., Dong, Y., Sumer, O., Bader, G. D. Cytoscape.js: A Graph Theory Library for Visualisation and Analysis. Bioinformatics 32 (2), 309–311 (2016).

20. Schloerke, B., Allen, J. plumber: An API Generator for R. R package version 1.3.3.9000, (2026). https://www.rplumber.io.

21. Tsugawa, H. et al. A lipidome landscape of aging in mice. Nat Aging 4, 709–726 (2024).

22. Mullen, T. D.; Hannun, Y. A.; Obeid, L. M. Ceramide Synthases at the Centre of Sphingolipid Metabolism and Biology. Biochem. J. 441 (3), 789–802 (2012).

22. Liu, Y., Xia, G., Zhu, S., Shi, Y., Huang, X., Wu, J., Xu, C., Du, A. Dijerential Transcriptomic Profiling of Lipid Metabolism and Collagen Remodeling in Fast- and Slow-Twitch Skeletal Muscles in Aging. FASEB J. 39 (2), e70335 (2025).

